# Colocalization of MOR1 and GAD67 in Mouse Nucleus Accumbens

**DOI:** 10.1101/574434

**Authors:** Chad Hinkle, Eduard Dedkov, Russell Buono, Thomas Ferraro

## Abstract

Current understanding of the rewarding and addictive effects of opioids involves mu-opioid receptor (MOR) binding within the nucleus accumbens (NAcc), a region of the ventral striatum. GABAergic neurotransmission in the NAcc potentiates the rewarding response to opioids, and in fact, drugs that stimulate GABAergic activity are also addictive, a phenomenon mediated in part by endogenous opioids. However, the neuroanatomical relationship between opioid and GABA systems is still unclear, and further study of the interaction between these neurotransmitter systems in the reward pathway is warranted. We report evidence supporting the direct interaction between GABAergic and opioidergic neurotransmitter systems within the mouse NAcc. Male and female FVB/NJ mice (12-16 months of age) were euthanized via carbon dioxide inhalation and brains processed for histology and immunohistochemistry (IHC). Coronal cryosections (10-12 um in thickness) were taken through the NAcc at the level of the anterior commissure. A mouse monoclonal antibody against GAD67, an enzyme catalyzing GABA production, was used in conjunction with an anti-mouse rhodamine red-X-labeled secondary antibody to identify GABAergic neurons in the NAcc. Alternating sections were stained for MOR using a fluorescein isothiocyanate (FITC)-conjugated rabbit polyclonal anti-MOR1 antibody. DAPI (4′,6-diamidino-2-phenylindole) was used to counterstain the nuclei. As expected, fluorescence microscopy results show that GAD67 staining is localized predominately in the neuronal cytoplasm, with approximately 40% of the population staining. The MOR1-FITC stain showed localization within the cytoplasm and plasma membrane as expected, however, there was also significant staining in the nuclear membrane and neuronal nucleus. MOR1-FITC was present in approximately 35% of the neuronal population. In separate experiments, we used double-immunostaining to study the co-expression of MOR1 and GAD67 within the same NAcc neurons. A similar localization pattern was detected with a small subset, approximately 2%, of neurons expressing both labels. There are few published reports of GAD67 and MOR1 co-expression within neurons of the NAcc. Previous studies of MOR expression show the receptor to be localized to the plasma membrane and, to a smaller degree, intracellularly. Here we found the MOR1 staining to be predominantly in the nucleus and nuclear membrane, with minimal expression on the plasma membrane. Further studies are in progress to validate the nuclear expression of MOR in GABAergic NAcc neurons. Results of the study suggest that a subset of mouse NAcc neurons express both MOR1 and GAD67, providing a direct intracellular link between opioid and GABAergic systems in the reward pathway. Preliminary intranuclear localization of MOR suggests a novel signaling pathway that may be important in fully elucidating neurobiological mechanisms underlying behaviors related to reward and addiction.

## INTRODUCTION

Two of the principal neurotransmitter systems that mediate reward signaling within the nucleus accumbens (NAcc), a neurochemically heterogeneous area of the brain implicated in reward processing and addiction, are opioids (endorphins, enkephalins) and GABA^12^. Opioid receptors comprise a family of multiple subtypes of G-protein coupled receptors (GCPRs), six and seven transmembrane-spanning receptors primarily located on the cell surface. Binding of exogenous or endogenous opioids, such as morphine or endorphins respectively, stimulates downstream signaling cascades to produce physiological effects^3,4^. Studies using agonists against the mu-opioid receptor (MOR) demonstrate that opioids modulate a broad range of nervous system functions, including sensation, cognition, emotion, and motor responses^5,6^. There has been extensive research aimed at uncovering how the opioidergic system elicits these wide variety effects. Specifically, the MOR has been shown to form heteromers with delta- and kappa-opioid subunits in addition to non-opioid GPCRs. Electron microscopy and *in situ* proximity ligation assay confirm that the close physical proximity of the MOR to other GPCRs intracellularly, such as serotonin, adrenergic, cannabinoid, and dopamine receptors, is in fact heteromerization^7,8^. Interacting with a variety of related and unrelated receptor families enhances MOR functional diversity and complexity. Multiple studies have confirmed the sparse distribution of the MOR within the rat NAcc using light microscopy and immunocytochemistry^4,6,9^. Furthermore, the interaction between the MOR and GABAergic neurons within the rat NAcc was investigated using electron microscopic immunocytochemical labeling with results demonstrating MOR immunoreactivity in the plasma membranes of GABAergic neurons, suggesting a functional link in the two systems^6^.

GABA is the major inhibitory neurotransmitter in the brain, found extensively on soma of projection and interneurons in all regions^6^. The neurotransmitter is synthesized via conversion of glutamate to GABA by glutamate decarboxylase (GAD) and is found in the majority of NAcc neurons^10,11,12,13^. The GABA_A_ receptor, an ionotropic chloride channel, and GABA_B_ receptor, an inhibitory GPCR, are expressed in the NAcc and have been implicated strongly in modulation of analgesia, anxiety, and depression^13^. Although the rewarding effects of opioid analgesic drugs, mediated via MOR, are thought to involve pathways of GABA signaling, the exact nature of the interaction between GABAergic and opioidergic systems in the NAcc is not fully established. Whereas cellular localization has been demonstrated between the opioidergic and GABAergic systems within cells of the rat NAcc^6^, we are not aware of such information in the brains of humans or mice.

In this study we used immunofluorescent staining to study the relationship between GABAergic and MOR-expressing neurons within the mouse NAcc. We provide direct evidence of the close proximity of the two neurochemical systems suggesting a functional relationship. Our results show MOR is localized to the nucleus and nuclear membrane of a subset of GABAergic neurons within the mouse NAcc. While the location of the receptor differs from the plasma membrane previously reported in the rat NAcc^6^, intracellular localization of opioid receptors within the golgi apparatus, endoplasmic reticulum, and nuclear membrane has been documented previously^14^. Another study demonstrated that endogenous peptides bind MOR on the plasma membrane and cause internalization to endosomes. Exogenous non-peptides, such as morphine, was found to distort this internalization by localizing to the golgi apparatus^15^. Our results suggest that endogenous opioidergic activity within the NAcc may modulate GABAergic signaling to produce its many behavioral effects via mechanisms that include binding of opioids to intracellular receptors.

## METHODS

Immunohistochemistry GAD67/ MOR1 colocalization. Male and female FVB/NJ mice (12-16 months of age) were euthanized by carbon dioxide inhalation followed by cervical dislocation. Brains were removed and immersion-fixed in 4% paraformaldehyde for 24 hours at 2-4°C. Brains were then washed in phosphate-buffered saline (PBS) for 24 hours (2-4°C), cryopreserved by incubating in a series of sucrose solutions of increasing concentration (0.25-1.5 M sucrose) for 1-1.5 hours each and embedded in Tissue-Tek^®^ OCT cryo-embedding compound. Frozen tissue was stored at −80°C until sectioning. Serial coronal sections were cut from each brain at a thickness of 10-12 um using a cryotome. Sections were placed on Superfrost^®^ Plus microscope slides three per slide and stored at −80°C until staining.

During tissue staining, every 4^th^ section was taken for cresyl violet (Nissl) staining. Nissl stains were performed in brain sections adjacent to those used for immunohistochemistry in order to examine cytoarchitecture and confirm regions of interest. As the NAcc can be difficult to distinguish from surrounding nuclei in the ventral striatum, Nissl staining allowed proper orientation of tissue sections and identification of related structures^5^. For Nissl staining, the sections were dehydrated in PBS for 10 minutes, rinsed in ddH_2_O for 1 minute and were then stained in a 0.1% solution of cresyl violet acetate for 20 minutes at room temperature. After staining, sections were dehydrated in a graded ethanol series, cleared in xylene and coverslipped using Permount.

Before double-immunostaining for GAD67 and MOR1 proteins, each antibody was individually validated by staining on separate brain tissue sections. Antibody characteristics are described in Table 1. For GAD67 immunostaining, sections were incubated in a mouse anti-GAD67 antibody at a dilution of 1:100 in PBS containing 1% Bovine Serum Albumin (BSA) for 30 minutes. To visualize primary antibody, sections were incubated in rhodamine red-X-conjugated goat anti-mouse secondary antibody (cat. 115-295-146; Jackson ImmunoResearch Lab., West Grove, PA) at a dilution of 1:200 in PBS. Slides were coverslipped with ProLong Gold Antifade Mounting media containing DAPI (cat. P36931; Molecular Probes, Inc., Eugene, OR) in order to counterstain the nuclei. For MOR1 immunostaining, brain sections were incubated in a rabbit anti-MOR1-FITC-conjugated antibody at a concentration of 1:20 in PBS containing 1% BSA for 30 minutes. Slides were coverslipped with ProLong Gold Antifade Mounting media containing DAPI. Negative controls for GAD67 and MOR1 antibodies were established by substituting primary antibodies with a PBS containing 1% BSA solution. For double-immunostaining, sections were first incubated in a primary antibody against GAD67 followed by incubation with a mixture of a rhodamine red-X-conjugated secondary antibody and a rabbit anti-MOR1-FITC conjugated antibody. Negative control was established by substituting primary antibody against GAD67 by using a PBS containing 1% BSA solution followed by incubation with a rhodamine red-X-conjugated secondary antibody only. Stained sections were observed under a Leica DM4000B LED fluorescent microscope (Leica Microsystem, Germany), ranging from 350-546/10-50nm excitation wavelengths and 460-585/40-50nm emission wavelengths, and images were captured onto a computer using Leica DFC360FX digital camera and Leica Software.

**TABLE 1.**
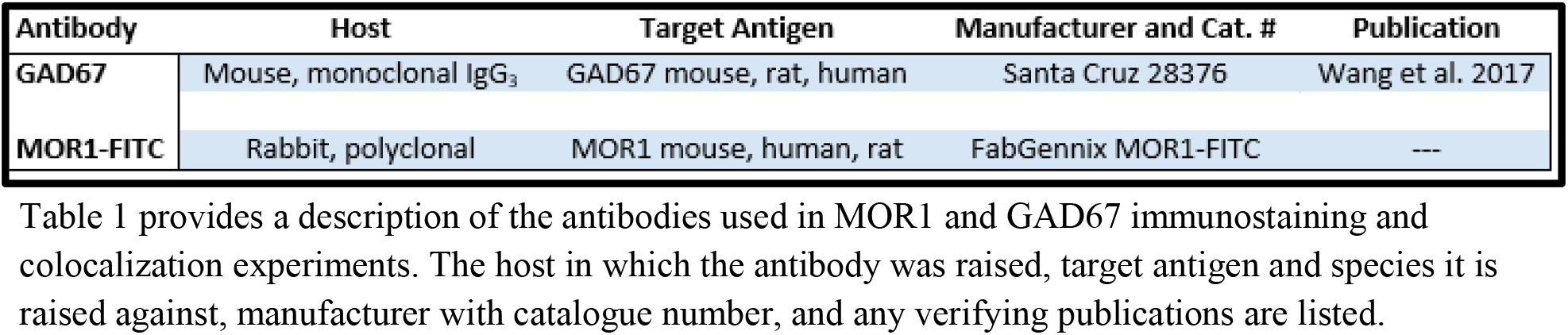
PRIMARY ANTIBODY PROFILES.

## RESULTS

### Colocalization of GAD67 and MOR1 in Mouse NAcc

Fluorescence immunohistochemistry was utilized to localize neurons containing GAD67 and MOR1 in the mouse NAcc. The ventral striatum, a region of the brain where the NAcc is found, is comprised of a population of neurons that are almost uniformly GABAergic^1,2^. Specifically, the NAcc efferent projections are believed to be GABAergic to the ventral tegmental area (VTA) and ventral pallidum (VP)^2^ and thus we expected a robust immunostaining of the GABA neurons in the NAcc. To identify GABAergic neurons, we used GAD67 as the neuronal marker^16,17^. Figure 1 (A,B,C) shows GAD67 immunostaining in the NAcc neurons using the primary mouse anti-GAD67 antibody visualized with a rhodamine red^TM^-X-conjugated secondary antibody. Nuclei were counterstained with DAPI. The distribution of GAD67 appeared to be primarily in the neuronal cytoplasm and nuclear membrane. The mu-opioid receptor (MOR) has been shown to be widely distributed in limbic and cortical regions such as the NAcc^18,19^. Therefore, we wanted to first demonstrate the presence of MOR in NAcc neurons of the mouse by targeting the most common MOR isoform, MOR1^20^. The anti-MOR1 antibody conjugated with FITC was used in conjunction with DAPI nuclear stain. Figure 1 (D,E,F) demonstrates MOR1 expression pattern in the NAcc neurons. Immunostaining of the MOR was abundant and appeared to be concentrated in the neuronal nucleus and nuclear membrane with minimal staining on the plasma membrane.

**Figure 1A-C.**
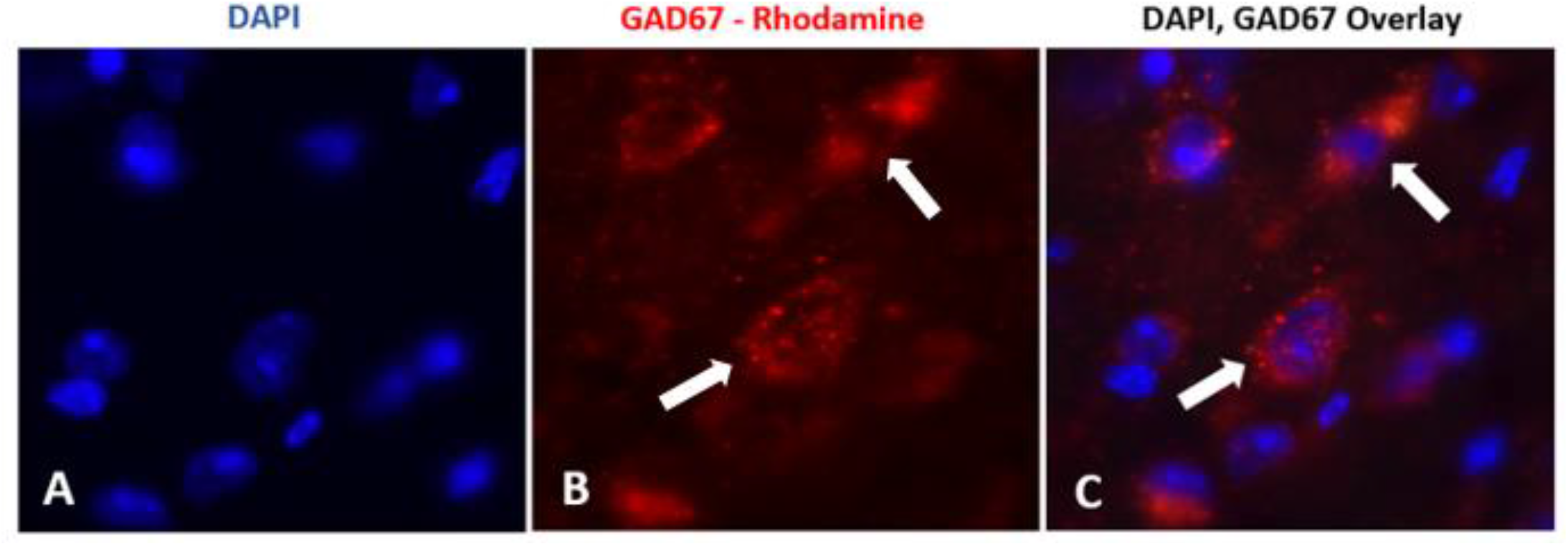
Immunostaining of mouse NAcc for glutamic acid decarboxylases 67 (GAD67). (A-C) Antibody detection of the GAD67 protein, as a marker for GABAergic neurons, is shown with the nuclear stain by DAPI. (A) Nuclear stain with DAPI visualized using a Leica A4 filter set (350/50nm excitation and 460/50nm emission wavelengths). (B) immunostaining with a mouse anti-GAD67 primary antibody visualized with a rhodamine red-X-conjugated secondary antibody using a Leica N2.1 filter set (546/10nm excitation and 585/40nm emission wavelengths). (C) overlay of DAPI and GAD67 staining. Note that in NAcc neurons GAD67 is predominantly present in cytoplasm (indicated by the white arrows). Images were taken at X40.

**Figure 1D-F.**
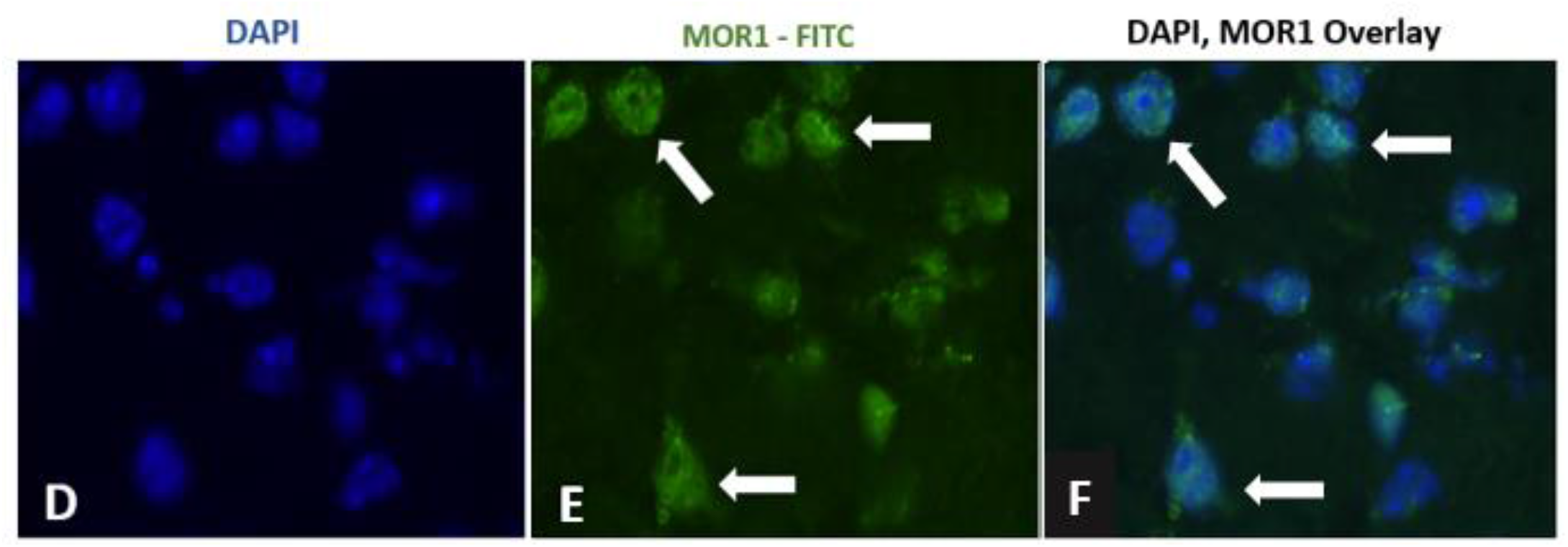
Immunostaining of mouse NAcc for Mu-type opioid receptor isoform 1 (MOR1). (D-F) Antibody detection of the MOR1 protein is shown with the nuclear stain by DAPI. (D) Nuclei stained by DAPI. (E) immunostaining with a rabbit anti-MOR1-FITC conjugated antibody visualized using a Leica L5 filter set (480/40nm excitation and 527/30nm emission wavelengths) and combined with a nuclear stain by DAPI. (F) abundant overlay of DAPI and MOR1 staining. Note that in NAcc neurons MOR1 protein expression appears to be predominantly localized inside the nucleus and on the nuclear membrane (indicated by the white arrows). Images were taken at X40.

After identifying neurons in the NAcc that express MOR1 and GAD67 individually, we wanted to determine whether there is a population of cells that co-express these proteins. A double-stain of GAD67 and MOR1 was performed using the anti-MOR1 antibody conjugated with FITC and the anti-GAD67 antibody visualized by rhodamine-red^TM^-conjugated secondary antibody. Figure 2 (A,B,C, and D) illustrates neurons in the NAcc co-expressing GAD67 and MOR1 targets. Based on the overlay of the red and green filters, we observed that the colocalized neurons express GAD67 in the cytoplasm while MOR1 tends to have a more intranuclear distribution. This distribution is similar to that observed in the single-label experiments for both of GAD67 and MOR1.

**Figure 2A-D.**
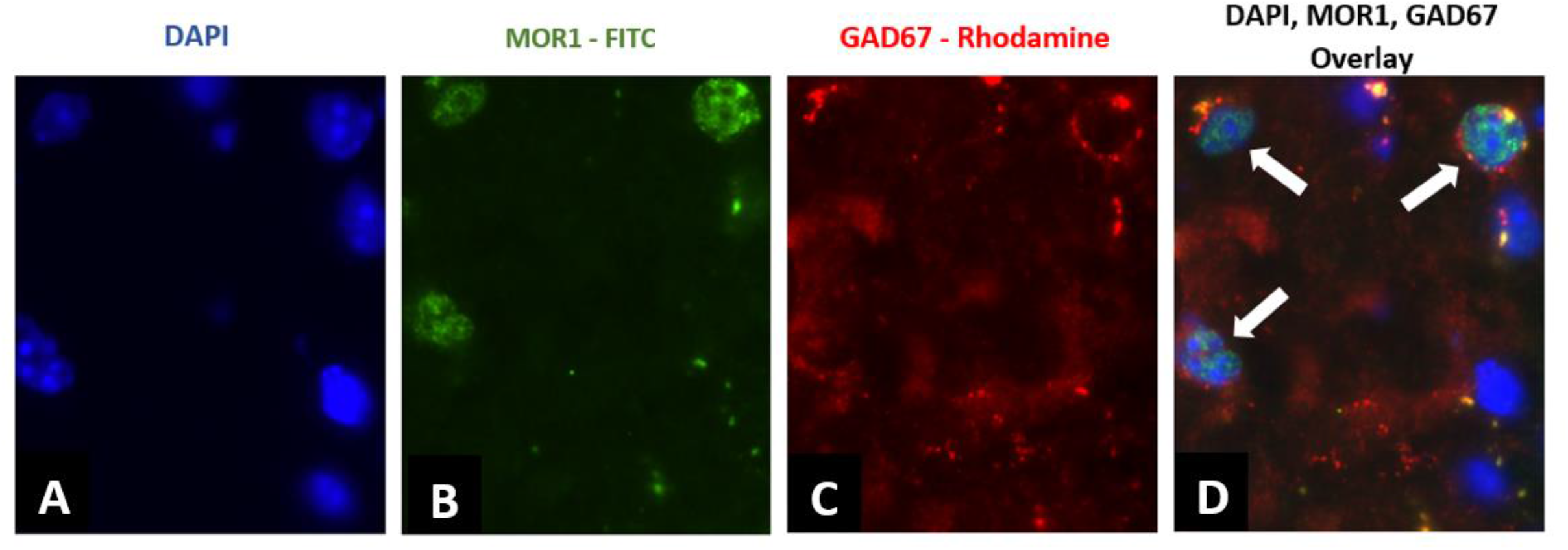
Concurrent localization of GAD67 and MOR1 proteins in mouse NAcc. (A-D) Double-immunostaining with a mouse anti-GAD67 and rabbit anti-MOR1 antibodies in the mouse NAcc combined with a DAPI nuclear stain. (A) Nuclear stain with DAPI visualized using a Leica A4 filter set. (B) immunostaining with a rabbit anti-MOR1-FITC conjugated antibody visualized using a Leica L5 filter set. (C) immunostaining with a mouse anti-GAD67 primary antibody visualized with a rhodamine red-X-conjugated secondary antibody using a Leica N2.1 filter set. (D) overlay of GAD67, MOR1, and DAPI staining demonstrates the co-localization of GAD67 and MOR1 proteins, in the cytoplasm and nuclei, respectively, of the NAcc neurons (indicated by the white arrows). Images were taken at X40.

## DISCUSSION

The results of the present study provide the first evidence for co-localization of MOR and GAD67 in cells within the mouse NAcc. While this has been documented previously in the rat, the subcellular distribution patterns were somewhat different^6^. Specifically, our study revealed three main findings. First, single-labeling of GABAergic neurons by GAD67 showed a clear cytoplasmic distribution without overlap of the nuclear stain. This is consistent with the literature as GABA expression in the cytoplasm of neuronal cell bodies is well documented^21,22^. We also found an abundance of GABAergic neurons within the NAcc which again is an expected finding based on literature reports^10,11,12,13,23^.

Second, we did find abundant MOR expression within the NAcc, which can be appreciated in Figure 1 (D-F). The MOR1-FITC stain was present in approximately 35% percent of the neuronal population in the NAcc. This is consistent with the literature which demonstrates that the NAcc has some of the highest levels of expression of opioid peptides in the CNS and should thus be abundant in staining^24,25,26,27,28,29^. A result that we did not expect was MOR expression appeared to be within the nuclear membrane. Demonstrated in Figure 1F, there was close overlap between the nuclear stain and MOR1 antibody in addition to cytoplasmic and plasma membrane staining. Current work is underway using confocal microscopy to confirm the subcellular localization of the MOR1 antibody. Other studies have shown that the MOR is found exclusively within the plasma membrane and cytoplasm^4,6,28^ While our findings show some plasma membrane and cytoplasm involvement, there have been reports that are consistent with our nuclear binding. The natural history of the opioid receptor has been hypothesized to begin with intracellular synthesis, followed by insertion into the plasma membrane with eventual internalization for destruction or ligand transport^14^. In a study of long-term memory formation, a nuclear role for opioids was proposed in the process of addiction and withdrawal^30^. It is unexpected that a purely synaptic process could lead to such a complicated chain of events without the involvement of gene alteration in the nucleus^30^. A recent study demonstrated that MOR activation induces internalization into endosomes when bound by endogenous peptides and MOR activation within golgi bodies with exogenous non-peptides^15^. In another study, antibodies against delta-opioid receptors were found to localize in the nucleus of neurohybrid cells, but mu- and kappa-receptors did not have specific binding in the nuclear preparations^14^. Another study used electron microscopy to identify radiolabeled MOR within the rat neostriatum; their results showed patches of MOR to be concentrated within neostriatal nuclear membranes^31^. Furthermore, studies involving the effect of opioids on cell growth postulate a nuclear involvement due to their inhibitor effect on cell division. Specifically, cerebellar neurons in 6-day old rats were shown to rapidly proliferate with the administration of naltrexone, an opioid antagonist^32^; a similar result was found in tumor cells after naltrexone exposure^33^. While the majority of MOR are known to be localized to the plasma membrane, our results and previous published studies support a nuclear involvement in opioid action. Early research on the behavioral effects of opiates postulated that they induce hyperpolarization of adjacent inhibitory interneurons, most likely GABA, resulting in disinhibition clinically^6,34,35^. More research is required to elucidate the functional correlation between MOR binding and GABA, but our results are consistent with this hypothesis of disinhibition.

A strength of this study is that we are the first to use immunohistochemistry to demonstrate the presence of MOR in GABAergic neurons of the mouse NAcc. Previous work using electron microscopy and immunocytochemistry demonstrates similar findings in rats^6^. When working with antibodies there is always concern for non-specific staining. While there are instances where we encountered staining of lipofuscin due to the age of the mice, we are confident our MOR and GABA overlap are true binding. A clue to distinguishing lipofuscin is its broad spectrum of auto-fluorescence^36^. The lipofuscin granules appeared very clearly between 480/40nm to 546/10nm excitation and 527/30nm to 585/40nm emission wavelengths, allowing them to be distinguished from true staining.

In conclusion, we have demonstrated that a subset of cells in the mouse NAcc express both GAD67 and MOR1, indicating a potential interaction between GABA and the mu-opioid receptor in opioid signaling within the same cell. Specifically, our results demonstrate the distribution of the MOR preferentially in the nucleus and nuclear membrane of GABAergic neurons. There has been little or no research showing the co-expression within the NAcc of mice, and here we add supporting data to the hypothesis that the effects of opioids may involve GABA signaling between and within neurons of the NAcc.

